# Cholinergic mu-opioid receptor deletion alters reward preference and aversion-resistance

**DOI:** 10.1101/2023.11.13.566881

**Authors:** Cambria R. Beane, Delainey G. Lewis, Nicolaus Bruns VI, Kat L. Pikus, Mary H. Durfee, Roman A. Zegarelli, Thomas W. Perry, Oscar Sandoval, Anna K. Radke

## Abstract

Heavy alcohol use and binge drinking are important contributors to alcohol use disorder (AUD). The endogenous opioid system has been implicated in alcohol consumption and preference in both humans and animals. The mu opioid receptor (MOR) is expressed on multiple cells in the striatum, however little is known about the contributions of specific MOR populations to alcohol drinking behaviors. The current study used mice with a genetic deletion of MOR in cholinergic cells (ChAT- Cre/Oprm1^fl/fl^) to examine the role of MORs expressed in cholinergic interneurons (CINs) in home cage self-administration paradigms. Male and female ChAT-Cre/Oprm1^fl/fl^ mice were generated and heterozygous Cre+ (knockout) and Cre- (control) mice were tested for alcohol and nicotine consumption. In Experiment 1, binge-like and quinine-resistant drinking was tested using 15% ethanol (EtOH) in a two-bottle, limited-access Drinking in the Dark paradigm. Experiment 2 involved a six-week intermittent access paradigm in which mice received 20% EtOH, nicotine, and then a combination of the two drugs.

Experiment 3 assessed locomotor activity, sucrose preference, and quinine sensitivity. Deleting MORs in cholinergic cells did not alter consumption of EtOH in Experiment 1 or 2. In Experiment 1, the MOR deletion resulted in greater consumption of quinine-adulterated EtOH in male Cre+ mice (vs. Cre-). In Experiment 2, Cre+ mice demonstrated a significantly lower preference for nicotine but did not differ from Cre- mice in nicotine or nicotine + EtOH consumption. Overall fluid consumption was also heightened in the Cre+ mice. In Experiment 3, Cre+ females were found to have greater locomotor activity and preference for sucrose vs. Cre- mice. These data suggest that cholinergic MORs are not required for EtOH, drinking behaviors but may contribute to aversion resistant EtOH drinking in a sex-dependent manner.

## 1. Introduction

Alcohol use disorder (AUD) is a significant public health crisis with 28.6 million adults in the United States ages 18 and older diagnosed with the disease (NIAAA, 2023). Cigarette use is twice as common in patients diagnosed with AUD vs. the general public (Weinberger et al., 2019). Binge drinking and heavy alcohol use contribute to the risk for developing AUD and are key symptoms of the disorder (NIAAA, 2023). Notably, compulsive drinking, defined as heavy drinking despite negative consequences, is an important characteristic of AUD (American Psychiatric Association, 2013). The prevalence of AUD and the resulting alcohol-related problems make research surrounding the neural underpinnings of this disease vital to developing novel and effective treatments and therapies.

The endogenous opioid system is known to be a critical regulator of alcohol consumption and preference in humans and animals (Gianoulakis, 2009; Méndez and Morales-Mulia, 2008). Increases in mu opioid receptor (MOR) expression and affinity have been associated with enhanced alcohol intake and dependence in rodents, non-human primates, and humans (Barr et al., 2007; Bart et al., 2005; Learn et al., 2001; Oslin et al., 2003). Genetic deletion of the mu opioid receptor (MOR) in mice reduces consumption (Becker et al., 2002; Hall et al., 2001; Roberts et al., 2000) as well as the acute stimulant and anxiolytic actions of ethanol (EtOH) (Ghozland et al., 2005). Further, systemic MOR antagonists decrease alcohol consumption in rodents (Gilpin et al., 2008; Heyser et al., 1999; Hyytiä and Kiianmaa, 2001) and the MOR antagonist naltrexone is one of three medications approved in the United States to treat AUD.

Despite this clear role of the endogenous opioid system and the MOR in alcohol drinking behaviors, the contributions of various MOR populations have yet to be fully identified. MORs are broadly expressed throughout the nervous system (Erbs et al., 2015), including in addiction-relevant nuclei such as the ventral tegmental area (VTA), dorsal striatum (DS), nucleus accumbens (NAc), and amygdala. Within each of these regions, MORs are expressed in multiple cell types. For example, in the striatum, MOR expression has been observed in both D1 and D2 subtypes of medium spiny neurons (MSNs) (i.e., direct and indirect pathway neurons) and in the cholinergic interneurons (CINs) (Cui et al., 2014; Jabourian et al., 2005; Oude Ophuis et al., 2014; Ponterio et al., 2013). MOR expression on D1-expressing, but not D2-expressing, MSNs contributes to morphine-induced locomotor sensitization (Severino et al., 2020). Furthermore, selective restoration of MOR expression in direct-pathway MSNs only partially restores opioid seeking behavior (Cui et al., 2014), suggesting that MOR-expressing neuronal subpopulations play distinct roles in behavior.

Although they comprise only 1-5% of striatal neurons, CINs are important regulators of dopamine (DA) release from terminals via nicotinic acetylcholine (ACh) receptor (nAChR) activation (Liu et al., 2022; Mohebi et al., 2023). MORs exhibit an inhibitory effect on striatal CIN excitability and striatal ACh release (Arttamangkul et al., 2021; Jabourian et al., 2005; Kiguchi et al., 2016; Ponterio et al., 2013), indirectly inhibiting DA overflow caused by a single-spike stimulus (Britt and McGehee, 2008; Yorgason et al., 2017). Thus, activation of CIN MORs could influence addictive behaviors by modulating DA signals associated with the response to rewarding stimuli such as drugs of abuse (Al-Hasani et al., 2021; Berlanga et al., 2003; Gonzales and Smith, 2015). Indeed, targeted deletion of MORs on cholinergic cells has been shown to selectively impair extinction of opioid-seeking (Severino et al., 2020). Additionally, alcohol exposure can activate CINs (Herring et al., 2004; Wadsworth et al., 2023) while both optogenetic and chemogenetic stimulation of NAc CINs has been shown to increase EtOH consumption (Kolpakova et al., 2022; Sharma et al., 2024).

Considering the role that striatal MORs are thought to play in regulating alcohol-drinking behaviors via alcohol-induced release of endogenous opioids (Heinz et al., 2005; Hermann et al., 2017; Lam et al., 2010; Méndez et al., 2010; Mitchell et al., 2012; Nealey et al., 2011; Olive et al., 2001; Perry and McNally, 2013; Richard and Fields, 2016; Zhang and Kelley, 2002), it is important to understand the relative contribution of MSN vs. CIN MOR expression to alcohol consumption. As such, we targeted CIN MORs by generating a transgenic line of mice with a deletion of the MOR on cholinergic cells and testing them across two models of alcohol consumption. In Experiment 1, we used a two-bottle, limited-access “drinking in the dark” (DID) paradigm (Sneddon et al., 2019) to model binge-and compulsive-like EtOH drinking. Three hours into the dark cycle, when mice are most active and likely to drink EtOH (Sprow and Thiele, 2012), they are given access to water and 15% EtOH in their home cages for a period of two hours. To model aversion-resistant drinking in this paradigm, we adulterated the EtOH with quinine, a bitter tastant (Hopf and Lesscher, 2014; Lesscher et al., 2010; Radke et al., 2021; Sneddon et al., 2019). Experiment 2 sought to further characterize alcohol drinking in these mice using a 24-h, two-bottle intermittent access (IA) paradigm. Because we saw no effect of MOR deletion on alcohol consumption in Experiment 1, and because acetylcholine release from CINs has been implicated in nicotine seeking (Leyrer-Jackson et al., 2021; Williams et al., 2017), mice in the IA experiment were also tested for nicotine consumption and preference. In this experiment, mice received EtOH, nicotine, and then a combination of the two drugs over a six-week period. Finally, we examined preference for sucrose, a non-drug reward, in Experiment 3.

## 2. Materials and Methods

### 2.1. Subjects

ChAT-Cre/Oprm1^fl/fl^ knockout mice (N = 123) were generated from breeding pairs purchased from the Jackson Laboratory (Bar Harbor, ME). Heterozygous ChAT-Cre males (B6.129S- Chattm1(cre)Lowl/MwarJ, strain #031661) were bred with homozygous *Oprm1* floxed females (B6;129- Oprm1tm1.1Cgrf/KffJ, strain #030074) to produce a Cre-mediated deletion of exons 2-3 of the MOR gene (*Oprm1*) on choline acetyltransferase (ChAT)-expressing neurons (**Figure 1A**). Following three crosses, the line was established and Cre- females were bred with Cre+ males to produce offspring that were Cre- or Cre+. All offspring were homozygous for floxed *Oprm1*.

**Figure 1.**
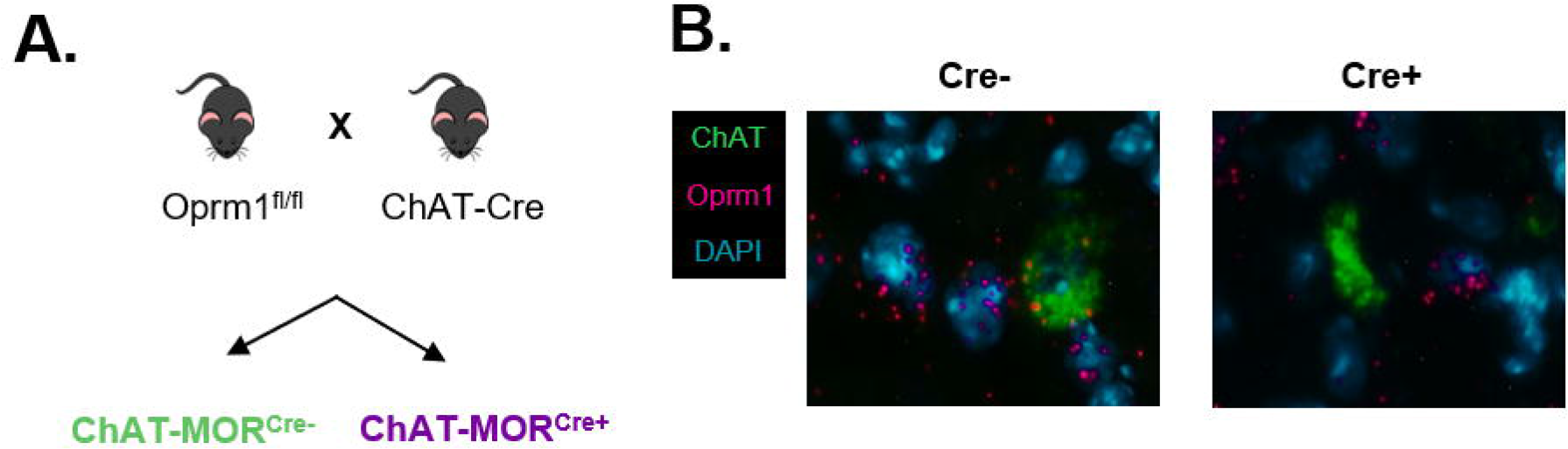
Knockout of mu opioid receptors on cholinergic neurons. A) Heterozygous ChAT-Cre males (B6.129S-Chattm1(cre)Lowl/MwarJ, strain #031661) were bred with homozygous *Oprm1* floxed females (B6;129-Oprm1tm1.1Cgrf/KffJ, strain #030074) to produce a Cre-mediated deletion of exons 2-3 of the MOR gene (*Oprm1*) on choline acetyltransferase (ChAT)-expressing neurons. ChAT-Cre/Oprm1^fl/fl^ Cre- offspring had intact mu opioid receptors (MOR) but Cre+ offspring were missing MORs on ChAT- expressing neurons only. **B)** Fluorescent *in situ* hybridization confirmed *Oprm1* deletion in *Chat* neurons only. Green = *Chat*, magenta = *Oprm1*, cyan = DAPI nuclear stain.

Group sizes were determined a priori using a power analysis on existing EtOH consumption data from our lab (using an alpha level of 0.05 with a specified level of power at 0.8), which demonstrated that a minimum of 14 mice were needed per group. Mice were housed at Miami University in standard shoe box rectangular mouse cages (18.4 cm x 29.2 cm x 12.7 cm) with food (LabDiet 5001) and reverse-osmosis (RO) water provided *ad libitum*. The cage lids were designed to allow for two test bottles to be present at the same time, with food placed between the water bottles. The animals resided in a temperature-controlled room with a 12:12 hour reverse light/dark cycle. For drinking experiments, mice habituated to the testing room and single housing conditions for at least one week before testing. All procedures were conducted in accordance with the Guide for the Care and Use of Laboratory Animals and were approved by the Miami University Institutional Animal Care and Use Committee.

### 2.2. Drugs

EtOH (15% or 20%) was made volume/volume in RO water. The concentrations of EtOH were chosen to facilitate comparison with our previous studies using DID (Sneddon et al., 2021, 2019) and IA (Radke et al., 2020; Perry et al., 2023). Quinine hemisulfate (Q1250-50G, Millipore-Sigma, St. Louis, MO) was prepared weight/volume in 15% EtOH or RO water. Nicotine tartrate salt (30 µg/mL) was prepared weight/volume in 20% EtOH solution or RO water. Nicotine concentrations are reported as free base. The concentration of nicotine used has previously been shown to support robust consumption in mice co-administering nicotine and EtOH (DeBaker et al., 2020). For DID, solutions were made fresh every day. For intermittent access, solutions were prepared on a weekly basis.

### 2.3. Fluorescent in situ hybridization

Validation of *Oprm1* deletion in the ChAT-Cre/Oprm1^fl/fl^ line was conducted using RNAScope (Advanced Cell Diagnostics, Newark, CA, USA) (**Figure 1B**). Brains from adult mice (8 Cre- and 8 Cre+) were extracted and flash frozen. A cryostat was used to collect sections (15 μm) containing the dorsal and ventral striatum. Sections were mounted on slides and fixed in chilled formalin and processed using probes for the cholinergic cell marker Mm-*Chat* (channel 1, cat#408731-C1) and the Mm-*Oprm1* gene (channel 2, cat#315841-C2). Slides were counterstained with DAPI and visualized using a Leica AX70 microscope.

### 2.4. Experiment 1: Effects of cholinergic MOR deletion on binge-like EtOH consumption and aversion- resistance

#### 2.4.1. EtOH drinking in the dark

Mice (15 male Cre-, 16 male Cre+, 15 female Cre-, and 17 female Cre+) self-administered EtOH in a two-bottle DID paradigm with access to EtOH for two hours a day, five days a week (Monday-Friday) for the duration of three weeks (**Figure 2A**). One of the bottles contained 15% EtOH, and the other bottle contained RO water. The side the bottles were presented on was alternated each day to prevent a side preference. Bottles were weighed before and after each drinking session and two “dummy” cages were positioned on the same rack as the mouse cages to account for any evaporation or spillage.

**Figure 2.**
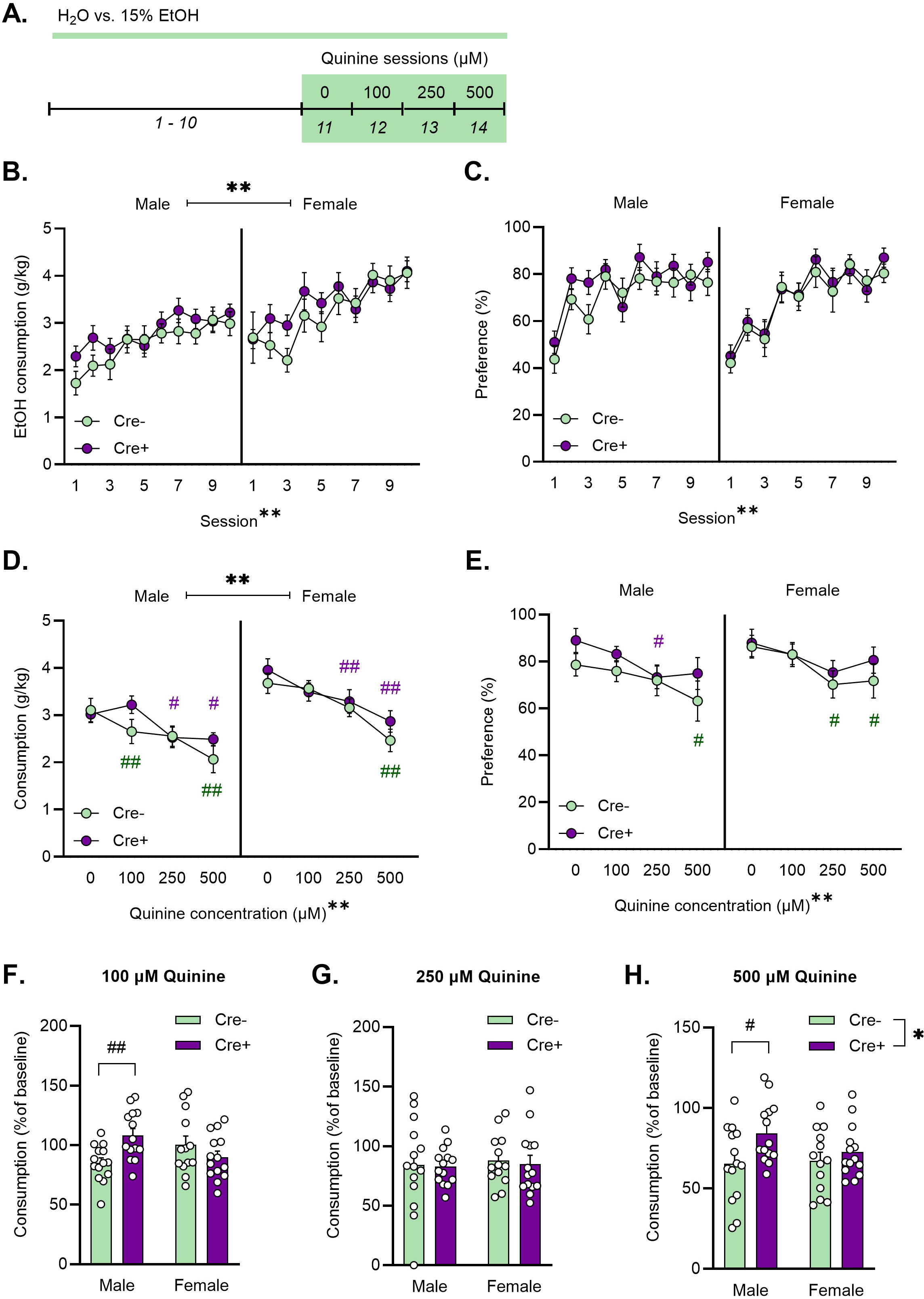
Cholinergic mu opioid receptor deletion does not affect ethanol consumption but does impact aversion-resistant drinking. A) Experimental timeline. Mice consumed 15% EtOH in a two-bottle choice DID task for 2 h/day over 10 sessions. Quinine hemisulfate was then added to the EtOH bottle in increasing concentrations over 4 sessions. **B)** Mice reliably drank 15% EtOH across sessions (** p < 0.01 main effect of session) and females consumed more than males (** p < 0.01 main effect of sex). MOR deletion did not lead to differences in 15% ethanol consumption or **C)** preference. **D)** Females drank more 15% ethanol with quinine than males (** p < 0.01 main effect of sex). and consumption decreased with increasing concentrations of quinine (** p < 0.01 main effect of concentration). The effects of quinine were dependent on sex, genotype, and quinine concentration (three-way interaction). # p < 0.05, ## p < 0.01 vs. 0-µm session (Dunnett’s test). **E)** Cholinergic MOR deletion did not affect preference during the quinine sessions although preference was lower in the presence of quinine (** p < 0.01 main effect of concentration). **F)** Cre+ (purple) males consumed more quinine-adulterated 15% EtOH than Cre- (green) males at the 100-µM concentration (calculated as a % of consumption on the 0- µM session) ( ## p < 0.01; Holm-Sidak test). **G)** Genotype did not affect consumption at the 250- µM concentration. **H)** Cre+ mice consumed more quinine adulterated 15% EtOH at the 500-µM concentration than Cre- mice (** p < 0.01 main effect of genotype) and this effect was most pronounced in males (# p < 0.05; Holm-Sidak test). There were no differences in the female mice at any concentration. N = 15 male Cre-, 16 male Cre+, 15 female Cre-, and 17 female Cre+.

During sessions 11-14, quinine hemisulfate was added to the 15% EtOH in increasing concentrations (0 µM, 100 µM, 250 µM, 500 µM) to test for aversion-resistant drinking.

#### 2.4.2. Water drinking in the dark

Following EtOH drinking, mice consumed water only for one week before testing of quinine sensitivity commenced (**Figure 3A**). The aversive effects of quinine in the absence of a drug were tested in two-hour drinking sessions identical to the previous sessions with EtOH. During these sessions, mice had access to two bottles, one with unadulterated RO water and the other with RO water and increasing concentrations of quinine presented over a series of 4 days (0 µM, 100 µM, 250 µM, 500 µM). Only 55 mice completed this portion of the experiment due to experimenter error.

**Figure 3.**
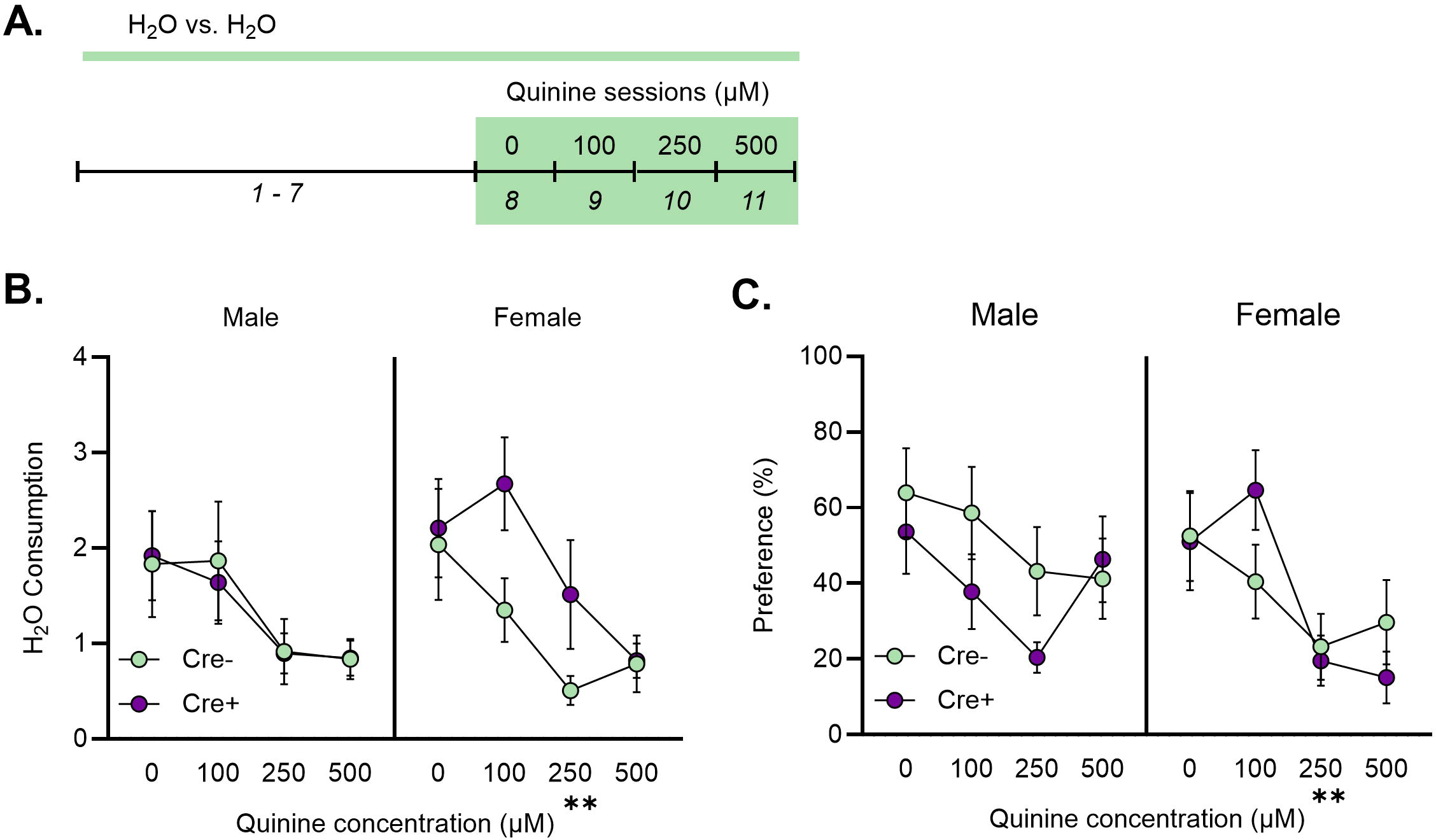
Cholinergic mu opioid receptor deletion does not affect quinine sensitivity. A) Experimental timeline. Mice consumed water for one week before being presented with increasing concentrations of quinine in a two-bottle choice task for 2 h/day. **B)** Consumption and **C)** preference for quinine-adulterated water decreased with increasing concentrations of quinine. ** p < 0.01 main effect (3- way ANOVA). N = 15 male Cre-, 16 male Cre+, 15 female Cre+, and 17 female Cre+.

### 2.5. Experiment 2: Effects of cholinergic MOR deletion on homecage self-administration of EtOH and nicotine

#### 2.5.1. Intermittent access to EtOH and nicotine

Mice were split into two cohorts roughly accounting for sex and genotype. Cohort one included 15 mice with 6 Cre+ mice (5 females and 1 male) and 9 Cre- mice (4 females and 5 males). Cohort two

included 15 mice with 9 Cre+ mice (6 females and 3 males) and 6 Cre- (2 females and 4 males). Treatment and care was identical between cohorts but varied in order. Mice self-administered 20% EtOH, 30 µg/mL nicotine, or EtOH and nicotine in combination in a 24-hour two-bottle intermittent access paradigm with access to the solutions every other day, three days a week (Mon/Wed/Fri) for the duration of two weeks per solution (**Figure 4A**). Drinking bottles were refilled on a daily basis 6 days/week at 12:30 p.m. On intervening sessions (T/Th), mice had access to two bottles of RO water. The bottle positions were alternated each day to prevent a side preference. Bottles were weighed before and after each drinking session and four “dummy” cages were positioned on the same rack as the mouse cages to track any evaporation or spillage. Animal body weights were recorded every day. Cohort one received 20% EtOH, followed by nicotine, and then the combined solution, each for two weeks. Cohort two began with the nicotine solution, followed by EtOH, and then the combined solution.

**Figure 4.**
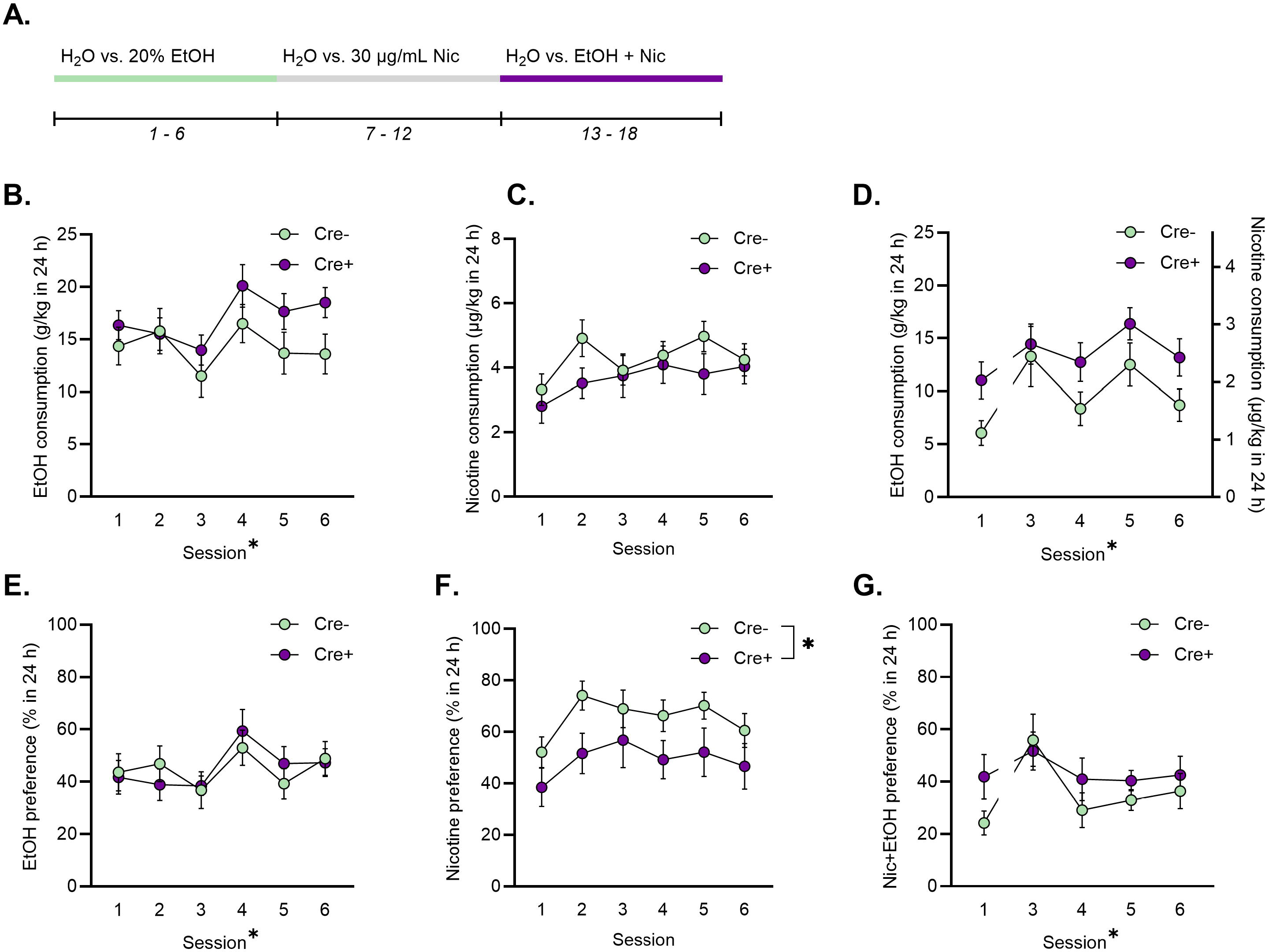
Cholinergic mu opioid receptor deletion decreased preference for nicotine during 24-hour intermittent access. Deletion of MOR did not impact **A)** Experimental timeline. Mice consumed 20% EtOH, 30 ug/mL nicotine, or a combination of EtOH and nicotine in a 24-h intermittent access task (3 sessions/week, 6 sessions per solution). The order of EtOH and nicotine presentation was reversed for half of the animals. **B)** 20% EtOH consumption, **C)** nicotine consumption, **D)** or 20% EtOH + nicotine consumption. **E)** Genotype did not impact 20% EtOH preference, **F)** decreased preference for nicotine, and **G)** did not affect 20% EtOH + nicotine preference. * p < 0.05 main effect (mixed effects analysis). N = 15 Cre+ and 15 Cre- mice.

**Figure 5.**
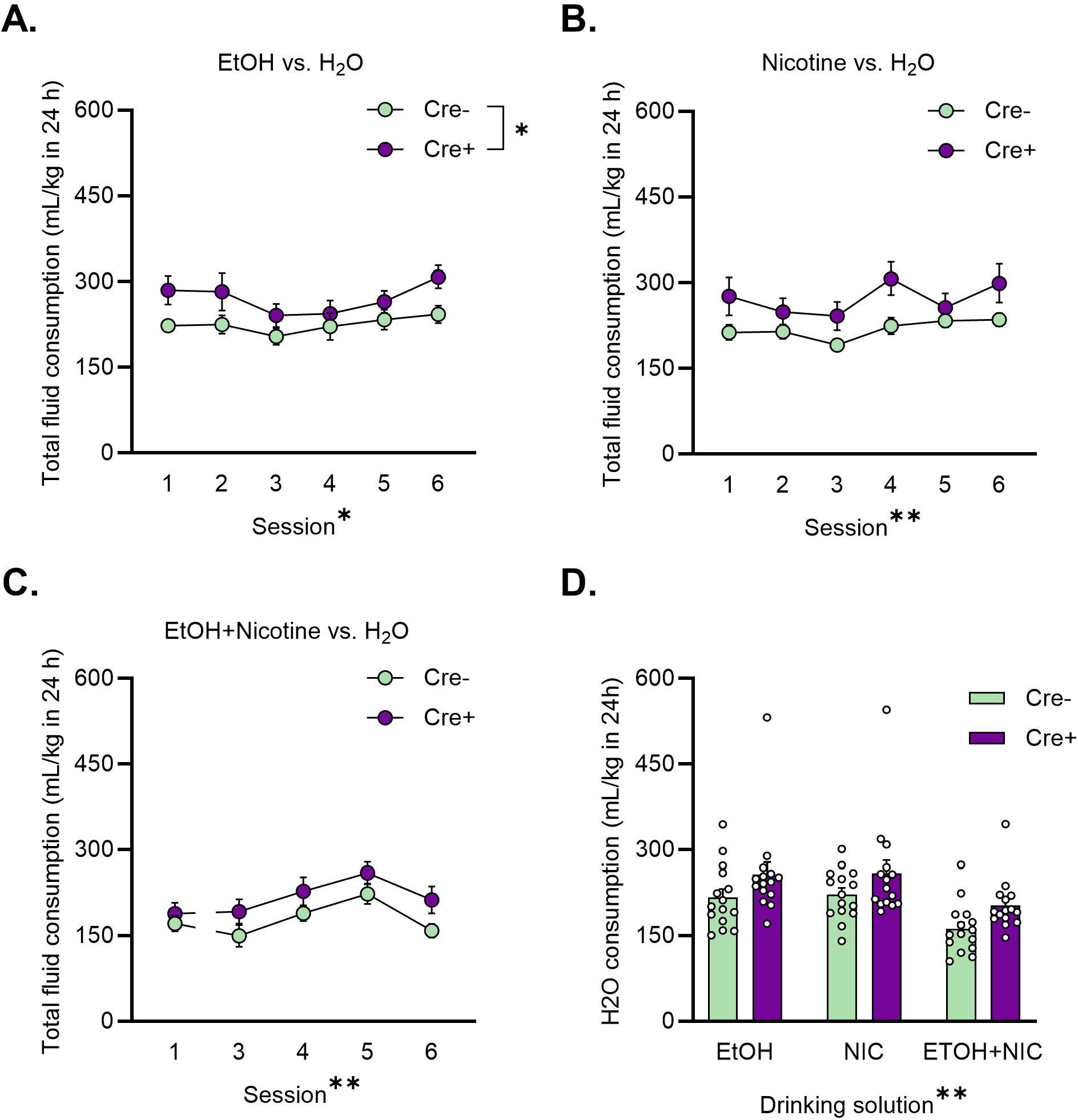
Cholinergic mu opioid receptor deletion increases fluid consumption. A) MOR deletion significantly increased total fluid consumption during EtOH sessions but not during **B)** nicotine or **C)** combined EtOH + nicotine sessions. **D)** There was a trend toward increased H_2_O in Cre+ mice across drinking solutions (p = 0.070). * p < 0.05, ** p < 0.01 main effect (2-way ANOVA). N = 15 Cre+ and 15 Cre- mice.

### 2.6. Experiment 3: Effects of cholinergic mu opioid receptor deletion on sucrose and quinine preference and locomotor behavior

#### 2.6.1. Locomotor test

To assess the effects of MOR deletion on baseline locomotion, mice (5 male, Cre-, 8 male Cre+, 7 female Cre-, and 10 female Cre+) were tested in activity chambers (14 in x 14 in x 8 in) that tracked movement via beam breaks (Omnitech Electronics, Inc, Columbus, OH). Mice were habituated to the procedure room for 30 minutes prior to recording behavior. Mice were tested for locomotion for just one session lasting 60 min in duration where activity was monitored and recorded.

#### 2.6.2. Sucrose and quinine preference test

One week after locomotor testing, mice (5 male, Cre-, 8 male Cre+, 7 female Cre-, and 10 female Cre+) self-administered sucrose in a continuous-access, two-bottle choice paradigm for five days (**Figure 7A**). One of the bottles contained a solution of sucrose and RO water with the concentration of sucrose increasing each day (0%, 0.1%, 0.5%, 1.0%, 5.0%), and the other bottle contained RO water. Following the sucrose preference test, the mice self-administered a quinine solution in the same two-bottle choice paradigm for five days. One of the bottles contained quinine added to RO water in increasing concentrations (0 µM, 25 µM, 100 µM, 250 µM, 500 µM), and the other bottle contained RO water. For both sucrose and quinine sessions, the side the bottles were presented on was alternated each day to prevent a side preference. Bottles were weighed before and after each drinking session and two “dummy” cages were positioned on the same rack as the mouse cages to account for any evaporation or spillage.

**Figure 6.**
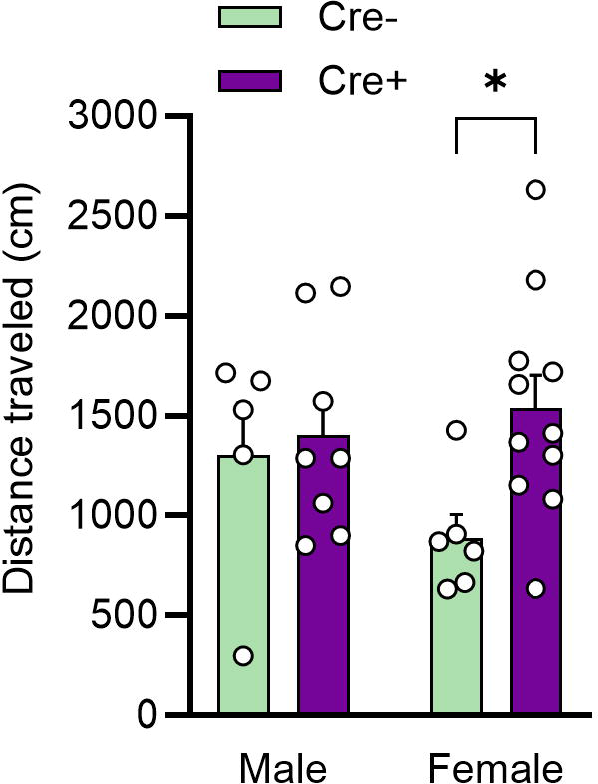
Cholinergic mu opioid receptor deletion increases locomotion in female mice. * p < 0.05 (Holm-Sidak test).

**Figure 7.**
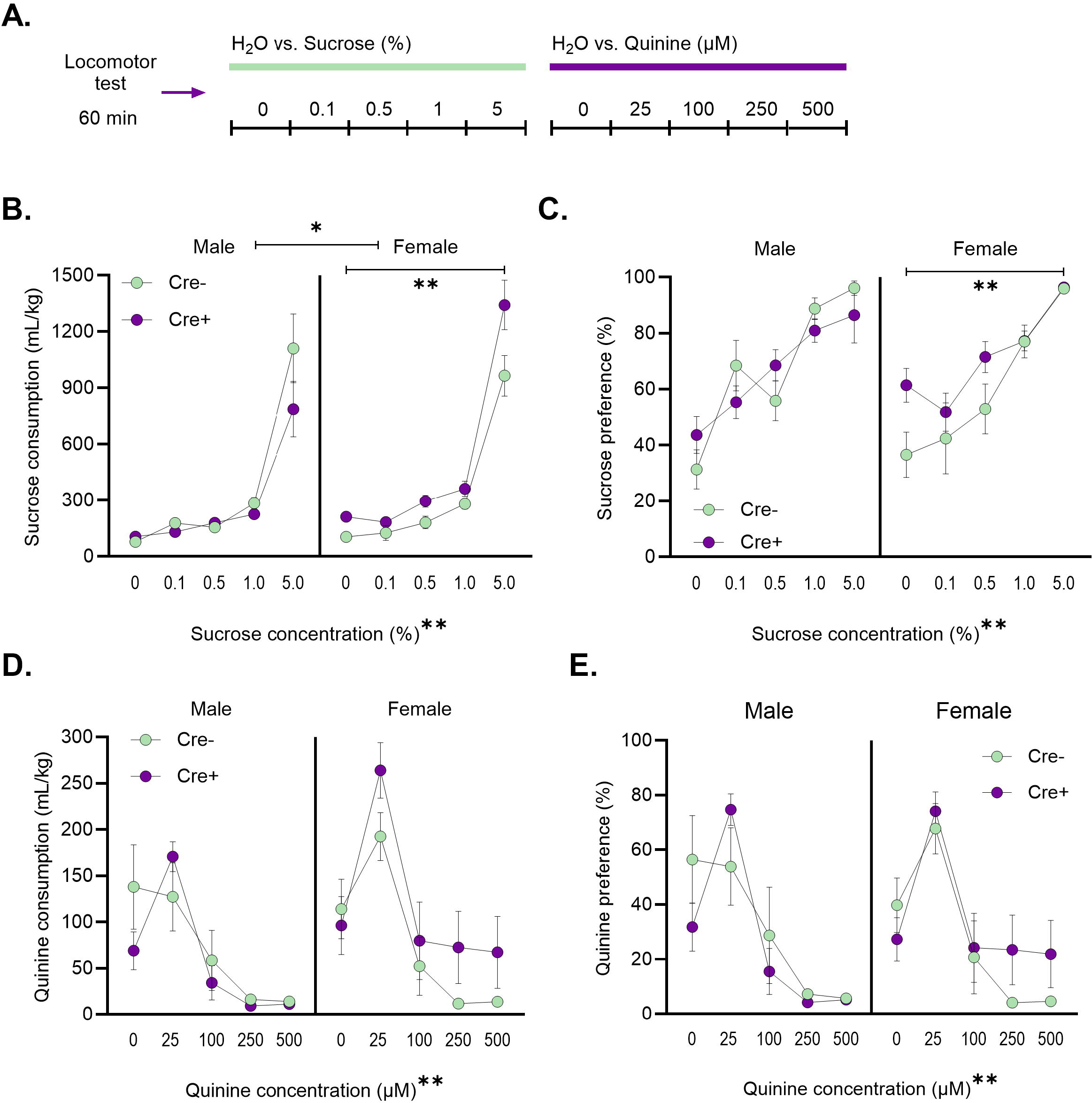
Cholinergic mu opioid receptor deletion increases sucrose consumption and preference in female mice. **A)** Experimental timeline. One week following a locomotion test, 24-h sucrose preference and sensitivity to quinine was tested for 5 sessions each. **B)** MOR deletion significantly increased sucrose consumption and **C)** preference in females. **D)** MOR did not impact quinine consumption or **E)** preference. *p < 0.05, **p < 0.01 main effect (3-way or 2-way ANOVA).

Solutions were available continuously except for when the bottles were weighed and refilled (approximately 1.5 h/day).

### 2.6. Data analysis

All analyses were performed using GraphPad Prism (v 10) software. Bottle weights were expressed as grams of EtOH, µ g of nicotine, or mL of water consumed per kg of bodyweight. The amount of solution consumed was measured by calculating *=(Initial Bottle Weight - Final Bottle Weight) - Average Dummy Bottle Weight*. Preference was measured by calculating *=(Volume of solution/Volume of solution + Water consumption)*100*. Total fluid consumption on test sessions and intervening sessions was calculated by summing consumption from the two bottles.

In all experiments, consumption (g/kg) and % preference were analyzed using three-way repeated measures analysis of variance (RM-ANOVA) and, in instances of missing data, mixed factor analyses with factors of genotype, sex, and session/concentration. Experiment 1 was fully powered to detect sex differences and sex was included as a factor in all analyses. Because sex did not interact with genotype when analyzing consumption or preference in Experiment 1 or in preliminary analyses for Experiment 2, the latter experiment was not fully powered to detect sex differences and data were collapsed across sex for final analyses and presentation. For Experiment 1, separate analyses were conducted for the EtOH/RO water, EtOH/quinine, and RO water/quinine sessions. Aversion-resistance was defined *a priori* as a change from baseline and *post-hoc* Dunnett’s tests were conducted to analyze changes in consumption based on sex and genotype during the quinine sessions. For Experiment 2, data were combined for each solution and separate analyses were conducted for the EtOH, nicotine, and combined solutions. Body weights and water consumption on intervening sessions were analyzed using 2-way ANOVA. In Experiment 3, initial analyses revealed sex-dependent effects, so the data are presented separately in each sex. Locomotor behavior was analyzed using 2-way ANOVA with sex and genotype as factors.

In Experiment 2, a technological error during the nicotine + EtOH portion of the study resulted in data from session 2 being lost so it is not included in the figures or analyses.

## 3. Results

### 3.1. Experiment 1: Cholinergic MOR deletion does not affect binge-like EtOH consumption

#### 3.1.1. Binge-like consumption

A 3-way RM-ANOVA yielded a main effect of both session (F (3.138, 185.2) = 18.080, p < 0.0001) and sex (F (1, 59) = 11.910, p = 0.001) for consumption by body weight (g/kg) (**Figure 2B**), demonstrating greater consumption in female ChAT-Cre/Oprm1^fl/fl^ mice vs. males. EtOH preference exhibited a significant main effect of session (F (3.654, 215.6) = 27, p < 0.0001) and a session x sex interaction (F (9, 531) = 2.320, p = 0.015) (**Figure 2C**). There were no main effects or interactions involving genotype for either measure. Consumption is calculated per body weight for each animal, so we analyzed body weights on the first day of the study (**Table 1**). A two-way ANOVA exhibited a main effect of sex (F(1, 59) = 76.05, p < 0.0001), but Cre+ and Cre- mice did not differ in weight.

**Table 1.**
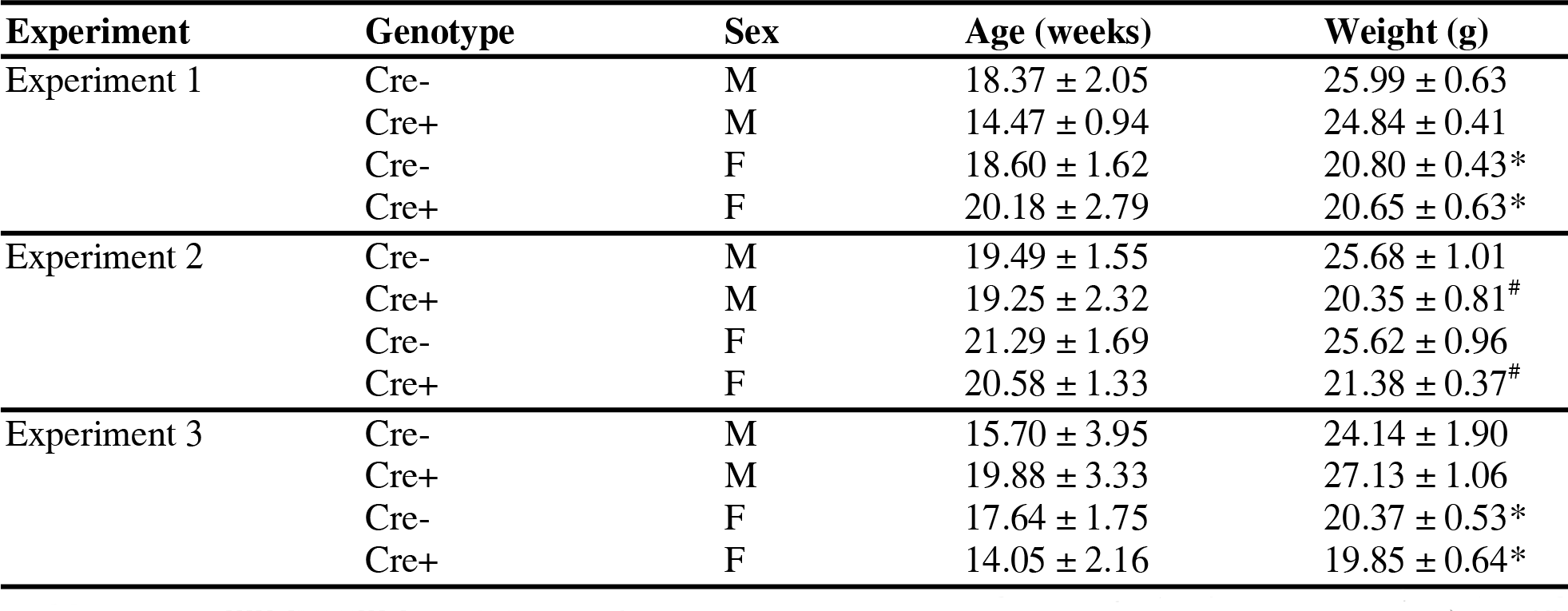
Age and weight of experimental animals at the start of each study. *p < 0.01 main effect of sex. # p < 0.01 main effect of genotype.

| Experiment | Genotype | Sex | Age (weeks) | Weight (g) |
| --- | --- | --- | --- | --- |
| Experiment 1 | Cre- | M | 18.37 ± 2.05 | 25.99 ± 0.63 |
|  | Cre+ | M | 14.47 ± 0.94 | 24.84 ± 0.41 |
|  | Cre- | F | 18.60 ± 1.62 | 20.80 ± 0.43* |
|  | Cre+ | F | 20.18 ± 2.79 | 20.65 ± 0.63* |
| Experiment 2 | Cre- | M | 19.49 ± 1.55 | 25.68 ± 1.01 |
|  | Cre+ | M | 19.25 ± 2.32 | 20.35 ± 0.81<sup>#</sup> |
|  | Cre- | F | 21.29 ± 1.69 | 25.62 ± 0.96 |
|  | Cre+ | F | 20.58 ± 1.33 | 21.38 ± 0.37<sup>#</sup> |
| Experiment 3 | Cre- | M | 15.70 ± 3.95 | 24.14 ± 1.90 |
|  | Cre+ | M | 19.88 ± 3.33 | 27.13 ± 1.06 |
|  | Cre- | F | 17.64 ± 1.75 | 20.37 ± 0.53* |
|  | Cre+ | F | 14.05 ± 2.16 | 19.85 ± 0.64* |

#### 3.1.2. Aversion-resistant drinking

For quinine-resistant drinking, 3-way RM-ANOVA revealed a main effect of quinine concentration on EtOH consumption (F (3, 153) = 35.670, p < 0.0001) (**Figure 2D**) and a significant main effect of sex (F (1, 51) = 9.299, p = 0.004). There was also a concentration x sex x genotype interaction (F (3, 153) = 2.933, p = 0.0354). To follow up on this interaction, separate 2-way RM ANOVAs were performed in male and female animals. In males, there was a significant main effect of concentration on consumption (F (3, 77) = 15.300, p < 0.0001) and a significant interaction between concentration and genotype (F (3, 77) = 3.938, p = 0.011). In females, there was a significant main effect of concentration (F (3, 75) = 21.130, p < 0.0001). Dunnett’s test were used to test aversion-resistance at each quinine concentration by comparing to the 0-µM session. 100-µM quinine reduced consumption in Cre- males (p = 0.037) only. For the 250-µM concentration, reductions were observed in Cre+ males (p = 0.022) and females (p = 0.006). Finally, consumption was reduced at the 500-µM quinine concentration in Cre- males and females (p < 0.0001 for both) and in Cre+ males (p = 0.013) and females (p < 0.0001).

A 3-way RM-ANOVA on EtOH preference during the quinine sessions found a significant main effect of session (F (3, 129) = 8.850, p < 0.0001) but no effects of sex or genotype (**Figure 2E**). Separate 2-way RM-ANOVAs found significant main effects of quinine concentration in male (F (3, 78) = 4.722, p = 0.004) and female (F (3, 75) = 5.747, p = 0.001) animals. There were no significant main effects or interactions with genotype. Preference for quinine-adulterated EtOH was significantly reduced in male Cre- mice at the 500-µM concentration (p = 0.037). In Cre+ males, preference was significantly reduced at the 250-µM concentration (p = 0.033) and the 500-µM concentration approached the threshold of significance (p = 0.063). In females, preference was significantly reduced in Cre- mice at the 250- (p =0.013) and 500-µM (p = 0.027) quinine concentrations. The Dunnett’s test approached the threshold of significance in female Cre+ mice at the 250-µM concentration (p = 0.054).

To further explore and visualize the 3-way interaction between concentration, sex and genotype found for EtOH consumption, g/kg intake on each quinine session (100, 250, and 500 µM) was expressed as a percent of consumption on the 0-µM session and 2-way ANOVAs were performed for each concentration. For the 100-µM session, there was a significant interaction between sex and genotype (F (1, 51) = 9.343, p = 0.004) and follow-up Holm-Sidak’s tests revealed a significant difference between Cre- and Cre+ consumption in male (p = 0.008) but not female (p = 0.185) mice (**Figure 2F**). No significant effects were observed at the 250-µM concentration (**Figure 2G**). At 500 µM, there was a significant main effect of genotype (F (1, 51) = 5.252, p = 0.026) and follow-up Holm-Sidak’s tests revealed a significant difference between Cre- and Cre+ consumption in male (p = 0.029) but not female (p = 0.710) mice (**Figure 2H**).

#### 3.1.3. Quinine sensitivity

Following the EtOH phase of the study, mice underwent a quinine sensitivity assessment during which they were exposed to increasing concentrations of quinine (0 µM, 100 µM, 250 µM, 500 µM) in water. A 3-way ANOVA found a significant main effect of quinine concentration on consumption (F (3, 152) = 11.070, p < 0.0001) but no main effect of sex (F (1, 51) = 0.270, p = 0.606) or genotype (F (1, 51) = 1.220, p = 0.275) (**Figure 3B**). A 3-way ANOVA of preference identified a main effect of quinine concentration (F (3, 153) = 10.510, p < 0.0001) but no main effect of sex (F (1, 51) = 1.394, p = 0.243) or genotype (F (1, 51) = 0.577, p = 0.451) (**Figure 3C**). The interaction between concentration, sex, and genotype approached the threshold for significance (F (3, 153) = 2.515, p = 0.061).

### 3.2. Experiment 2: Cholinergic MOR deletion reduces nicotine preference

ChAT-Cre/Oprm1^fl/fl^ mice were next tested for EtOH and nicotine consumption using the Intermittent Access model. As with binge-like consumption in Experiment 1, there were no effects of MOR deletion on EtOH consumption, although a 2-way mixed effects model revealed a main effect of session (F (3.414, 94.900) = 3.601, p = 0.013) (**Figure 4B**). There were similarly no effects of genotype on consumption of nicotine or the nicotine + EtOH solution when analyzed using 2-way RM-ANOVA (**Figure 4C-D**). The main effect of session approached the threshold for significance for the nicotine solution (F (3.532, 98.890) = 2.252, p = 0.077) and was significant for nicotine + EtOH (F (2.924, 81.860) = 6.639, p = 0.001).

A 2-way mixed effects model on 24-hour EtOH preference revealed a main effect of session (F (3.405, 94.650) = 2.928 p = 0.032) but no effect of genotype (**Figure 4E**). Preference for the nicotine solution was significantly lower in Cre+ mice with MOR deletion, as suggested by a main effect of genotype (F (1, 28) = 7.347, p = .0113) (**Figure 4F**). 2-way ANOVA found no significant main effect of session or interaction for this solution. Finally, for the combined solution, preference was not affected by genotype but did vary with session (F (3.028, 84.790) = 4.544, p = .005) (**Figure 4G**).

Effects of genotype on nicotine preference but not consumption suggested that the MOR deletion in cholinergic cells was influencing consumption from the H_2_O bottle. Analyses of total fluid and H_2_O consumption suggested increased H_2_O intake in Cre+ mice. For EtOH, a 2-way mixed effects analysis of total fluid consumption revealed main effects of session (F (3.680, 103.000) = 3.174, p = 0.020) and genotype (F (1, 28) = 4.370, p = 0.046) but no interaction (**Figure 5A**). There was a main effect of session for the nicotine solution (F (2.618, 72.250) = 5.233, p = 0.004) and the combined nicotine + EtOH solution (F (2.589, 72.490) = 8.678, p < 0.001) (**Figure 5B-C**). The main effect of genotype approached the threshold of significance for both nicotine (F (1, 28) = 3.990, p = 0.056) and the combined solution (F (1, 28) = 3.553, p = 0.070). A 2-way ANOVA of H_2_O consumption on intervening days (i.e., T/Th) found a main effect of solution (F (1.533, 42.920) = 35.350, p < 0.0001) and the main effect of genotype approached the threshold for significance (F (1, 28) = 3.553, p = 0.070) (**Figure 5D**). Because consumption is calculated as a function of the animal’s bodyweight, we also analyzed bodyweights on the first day of the study and found that Cre+ mice weighed less than Cre- mice (main effect of genotype (F(1, 26) = 31.19, p <0.0001) (**Table 1**).

### 3.3. Experiment 3: Cholinergic mu opioid receptor deletion increases sucrose preference

#### 3.3.2. Locomotion

*M*ice were first tested for locomotor activity in a 60 min session (**Figure 6**). A 2-way ANOVA revealed a main effect of genotype that approached the threshold for significance (F (1, 26) = 3.810, p = 0.062). Post-hoc Holm-Sidak’s tests revealed a significant difference between Cre- and Cre+ females (p = 0.034) but not males (p = 0.734).

#### 3.3.1. Sucrose and quinine preference test

Sucrose consumption was influenced by genotype in a sex-dependent manner (**Figure 7B**). A 3- way RM-ANOVA on sucrose consumption found significant main effects of concentration (F (4, 104) = 132.5, p < 0.0001) and sex (F (1, 26) = 4.992, p = 0.034). There was also a sex x genotype interaction (F (1, 26) = 9.398, p = 0.005) and a concentration x sex x genotype interaction (F (4, 104) = 3.927, p = 0.005). To follow up on these interactions, separate 2-way RM ANOVAs were performed in male and female animals. In males, there was only a significant main effect of sucrose concentration (F (1.112, 12.230) = 45.190, p < 0.0001). In females, there was a significant main effect of sucrose concentration (F (1.313, 19.700) = 96.510, p < 0.0001) and genotype (F (1, 15) = 9.586, p = 0.007), demonstrating greater sucrose consumption in Cre+ (vs. Cre-) females.

Sucrose preference was also analyzed using a 3-way RM-ANOVA (**Figure 7C**), which revealed a significant main effect of sucrose concentration (F (2.435, 63.31) = 33.050, p < 0.0001). There was also a sucrose concentration x genotype interaction (F (4, 104) = 2.569, p = 0.042) and a sucrose concentration x sex x genotype interaction (F (4, 104) = 0.208, p = 0.934). Separate 2-way RM ANOVAs were performed in male and female animals to follow up on these interactions. In males, there was only a significant main effect of sucrose concentration (F (4, 55) = 18.670, p < 0.0001). In females, there was a significant main effect of sucrose concentration (F (4, 75) = 20.140, p < 0.0001) and genotype (F (1, 75) = 6.781, p = 0.011).

Sensitivity to quinine was not influenced by sex or genotype (**Figure 7D-E**). Quinine consumption was analyzed using a 3-way RM ANOVA, revealing a significant main effect of quinine concentration (F (4, 104) = 31.82, p < 0.0001) and no other significant main effects or interactions. For quinine preference, a 3-way RM ANOVA found only a significant main effect of quinine concentration (F (2.320, 60.33) = 30.440, p < 0.0001).

## 4. Discussion

Here, we investigated the effects of the deletion of cholinergic MORs on nicotine consumption and binge- and compulsive-like EtOH consumption. Male and female ChAT-Cre/Oprm1^fl/fl^ mice demonstrated robust levels of alcohol consumption in the DID task, with females consuming more than males. These levels of consumption are similar to those typically observed in C57BL/6J mice using the same parameters (Schuh et al., 2022; Sneddon et al., 2021, 2019). However, Cre+ mice with MOR deletion did not differ in their consumption or preference for 15% EtOH. Similarly, ChAT-Cre/Oprm1^fl/fl^ mice consumed 20% EtOH readily in the intermittent access experiment but there were no differences between genotypes. When testing aversion-resistant drinking, we observed that Cre+ male mice demonstrated a tendency to consume more quinine-adulterated 15% EtOH, specifically at the 100-µM and 500-µM concentrations. The deletion of cholinergic MORs did not alter 24-h consumption of the nicotine or nicotine + 15% EtOH solutions, but Cre+ mice did exhibit a lower preference for nicotine than Cre- mice. Finally, we observed greater consumption of sucrose and locomotor activity in Cre+ females. Overall, these findings demonstrate that cholinergic MORs are not required for EtOH self-administration behaviors and suggest that CINs may be involved in aversion resistance.

Acetylcholine signaling has an established role in mediating alcohol reinforcement. This relationship is demonstrated by the ability of nicotine to alter EtOH self-administration in animal models and alcohol consumption in humans, although the direction of the effect is variable and dependent on a number of experimental factors (for review see Hendrickson et al., 2013). EtOH consumption is also reduced by the nAChR partial agonist varenicline (Blomqvist et al., 1996; Ericson et al., 2009; Hendrickson et al., 2010; Steensland et al., 2007; Wouda et al., 2011), and these effects rely at least in part on cholinergic regulation of dopamine release in the NAc core (Feduccia et al., 2014; Nadal et al., 1998; Schilaty et al., 2014). CINs are the primary source of ACh in the striatum and ablation of these neurons reduced EtOH-induced increases in NAc dopamine (Loftén et al., 2023, 2021). Targeted excitation of NAc CINs has also been shown to increase EtOH consumption in mice (Kolpakova et al., 2022). Although the cholinergic system clearly plays an important role in mediating alcohol drinking behaviors, our results in ChAT-Cre/Oprm1^fl/fl^ mice demonstrate that cholinergic MOR expression is not a necessary component of this process. Thus, endogenous opioid release in the striatum following alcohol exposure likely exerts its behavioral effects through MORs expressed on D1 and/or D2 receptor-expressing MSNs or through delta opioid receptors (DOR), which are also found on CINs (Arttamangkul et al., 2021; Le Moine et al., 1994). The latter possibility is supported by findings demonstrating that a DOR, but not MOR, agonist stimulated EtOH intake when infused into the NAc (Barson et al., 2009) and that DOR antagonism in the NAc prevents EtOH-induced release of DA (Acquas et al., 1993).

Although cholinergic MOR deletion did not affect EtOH consumption or preference in the limited access DID or intermittent access tasks, intriguing sex-dependent effects were observed on quinine-resistant drinking. At both the lowest and highest concentrations of quinine, mice with MOR deletion were more resistant to aversion than their Cre- counterparts and this effect was stronger in males. Put another way, Cre- male controls were sensitive and responded similarly to quinine at the 100 µM and 250 µM concentrations while mice with MOR deletion only reduced responding at the highest (250 and 500 µM) concentrations. It is unclear at present whether sex differences in this effect represent sex differences in the cholinergic system or a ceiling effect resulting from higher consumption in female mice. Because CINs are known to mediate other types of behavioral flexibility (Aoki et al., 2018, 2015; Okada et al., 2018), including following chronic alcohol exposure (Galaj et al., 2019; Ma et al., 2022), it is interesting to speculate that functional differences in CINs induced by the MOR deletion resulted in a reduced capacity for flexibility in ChAT-Cre/Oprm1^fl/fl^ Cre+ mice. The finding that CINs may participate in aversion-resistant alcohol drinking is also interesting in light of studies demonstrating a role for NAc core neurons in quinine-resistant consumption in rats and mice (Seif et al., 2013; Sneddon et al., 2021). After demonstrating that global inhibition of NAc core neurons reduces aversion-resistant drinking while targeted inhibition of D1 and D2 receptor-expressing neurons in this region does not, we previously speculated that CINs may be important regulators of this behavior (Sneddon et al., 2021).

Similar to EtOH, nicotine induces ACh release from striatal CINs (Mansvelder et al., 2003; Rada et al., 2001) and endogenous opioid release in the striatum (Dhatt et al., 1995; Houdi et al., 1991; Marty et al., 1985). Indeed, multiple behavioral and cellular effects of nicotine are absent in transgenic mice missing MOR or endogenous opioid peptides (Berrendero et al., 2010; Trigo et al., 2009; Walters et al., 2005; Yoo et al., 2004). Further, inhibition of CINs was shown to reduce cue-induced reinstatement of nicotine seeking (Leyrer-Jackson et al., 2021). Based on these lines of evidence, we hypothesized that similar results would be seen for nicotine and EtOH drinking in mice with cholinergic MOR deletion.

Instead, we observed that nicotine preference, but not consumption, was attenuated in Cre+ vs. Cre- mice. Assessing two-bottle preference for drugs such as nicotine is a common approach because it reflects reinforcement (Collins et al., 2012; Pogun et al., 2017), but the measure can be confounded by factors such as total fluid consumption. The effect on nicotine preference seen here was likely more robust than for consumption because Cre+ mice also tended to drink more water (resulting in greater total fluid consumption) across all phases of the experiment. A small decrease in nicotine consumption on top of increased water intake can explain the significant result with nicotine preference. Still, it is notable that this effect was not observed during the EtOH drinking sessions. Whether resulting directly from the lack of MORs or adaptations in the cholinergic system brought on by the genetic manipulation (e.g., alterations in CIN excitability or nAChR expression), the discrepant results between EtOH and nicotine preference suggest that the cholinergic system plays a stronger role in nicotine than EtOH drinking. It will be important to confirm these conclusions regarding nicotine vs. EtOH preference in future experiments using conditioned place preference or other choice procedures. We also cannot completely rule out the possibility that the order of presentation for EtOH and nicotine affected the results, due to the small sample size in each cohort.

To determine whether our results could be explained by general increases in locomotion or sensitivity to non-drug rewards, we also assessed locomotor activity and sucrose preference in ChAT- Cre/Oprm1^fl/fl^ mice. Cholinergic MOR deletion increased locomotion and preference for sucrose in female mice. Differences in locomotion or interaction with the drinking bottles could influence the consumption and preference measures used in our study. For example, Cre+ mice tended to consume more fluids than controls and had lower bodyweights in the intermittent access experiment. The combination of greater locomotor activity, higher fluid consumption, and lower bodyweights suggest a pattern of hyperactivity in the Cre+ ChAT-Cre/Oprm1^fl/fl^ mice that could have confounded some of the results presented here. It will be important to perform additional characterization of ChAT-Cre/Oprm1^fl/fl^ line to address these issues.

While the current results support the conclusion that cholinergic MORs are not involved in EtOH consumption or preference, there are some caveats to the approach. First, MOR deletion was targeted to all cholinergic cells, including the ChAT+ neurons clustered in the pons and basal forebrain. It is thought that cholinergic neurons in the pedunculopontine tegmentum (PPTg) express MOR (Serafin et al., 1990) and nuclei in the pons provide cholinergic input to the VTA and striatum that influences drug-induced reward and locomotion (Corrigall et al., 1999; Steidl et al., 2017). As such, we cannot isolate the impacts of the current genetic deletion to striatal CINs. Additionally, genetic deletion of MOR was not restricted in time and we have not investigated how this manipulation alters functional properties of cholinergic neurons. It is possible that the observed effects are the result of adaptations in cholinergic neurons or in downstream circuits. Such compensatory adaptations could also mask the function of MORs in behavior, meaning that, despite observing negative effects, we also cannot completely rule out a role for cholinergic MORs in EtOH drinking. Another point of interest is that CIN MOR expression fluctuates significantly throughout the day (Jabourian et al., 2005), which may be responsible for diurnal rhythms in CIN regulation of dopamine release and conditioned responses to reward-paired cues (Stowe et al., 2022). The role of cholinergic MORs in EtOH and nicotine drinking behaviors may consequently be time dependent and not fully captured by the methods used here. We also did not take blood samples from our animals and thus do not have any information on how the reported levels of consumption correspond to levels of EtOH or nicotine in the blood for the ChAT-Cre/Oprm1^fl/fl^ mice.

## Conclusions

Deletion of *Oprm1* on cholinergic neurons did not alter EtOH consumption or preference using two models of alcohol intake but nicotine preference was reduced in mice with the deletion. Quinine-resistant EtOH consumption was greater in male mice following cholinergic MOR deletion while preference for sucrose was greater in females with the deletion. These results suggest that cholinergic MORs participate in preference for rewarding substances. Further, while they are not required for consumption of alcohol alone, cholinergic MORs may influence the tendency to drink despite negative consequences. Considering these promising results, future studies examining long-term self-administration and corresponding changes in neurophysiology, neurochemistry, and gene expression induced by cholinergic MOR deletion or CIN manipulations will be well justified.

## Funding sources

This research was supported by NIH grants R15 AA027915 (AKR), awards from the Office of Research for Undergraduates (CRB, NB, DGL, KLP, MHD) at Miami University, and the Broadening Undergraduate Research Participation Program in the Department of Psychology at Miami University.

**Conflict of interests:** The authors declare no conflicting interests in this work.

**Abbreviations:** AUD: Alcohol Use Disorder; MOR: mu opioid receptor; EtOH: ethanol; VTA ventral tegmental area; DS: dorsal striatum; NAc: nucleus accumbens; MSN: medium spiny neuron; CIN: cholinergic interneuron; DA: dopamine; nAChR: nicotinic acetylcholine receptor; DID: drinking in the dark; ChAT: choline acetyltransferase; IA: intermittent access

## Acknowledgements

The authors are grateful to Ezra Eccles and Harrison Carvour for technical assistance.

## Author contributions

Cambria Beane: Conceptualization, Formal Analysis, Investigation, Writing - Original Draft, Visualization. Delainey Lewis: Conceptualization, Formal Analysis, Investigation, Writing - Original Draft. Nicolaus Bruns VI: Conceptualization, Formal Analysis, Investigation. Kat Pikus: Conceptualization, Investigation, Writing – Original Draft. Mary Durfee: Conceptualization, Investigation. Roman Zegarelli: Investigation, Project Administration. Thomas Perry: Investigation, Visualization. Oscar Sandoval: Investigation, Formal Analysis, Writing – Original Draft. Anna Radke: Conceptualization, Formal Analysis, Writing - Review and Editing, Visualization, Supervision, Funding Acquisition.

## References

1. Acquas, E., Meloni, M., Di Chiara, G., 1993. Blockade of delta-opioid receptors in the nucleus accumbens prevents ethanol-induced stimulation of dopamine release. Eur. J. Pharmacol. 230, 239– 241.

2. Al-Hasani, R., Gowrishankar, R., Schmitz, G.P., Pedersen, C.E., Marcus, D.J., Shirley, S.E., Hobbs, T.E., Elerding, A.J., Renaud, S.J., Jing, M., Li, Y., Alvarez, V.A., Lemos, J.C., Bruchas, M.R.., 2021. Ventral tegmental area GABAergic inhibition of cholinergic interneurons in the ventral nucleus accumbens shell promotes reward reinforcement. Nat. Neurosci. 24, 1414–1428.

3. Aoki, S., Liu, A.W., Akamine, Y., Zucca, A., Zucca, S., Wickens, J.R., 2018. Cholinergic interneurons in the rat striatum modulate substitution of habits. Eur. J. Neurosci. 47, 1194–1205.

4. Aoki, S., Liu, A.W., Zucca, A., Zucca, S., Wickens, J.R., 2015. Role of Striatal Cholinergic Interneurons in Set-Shifting in the Rat. J. Neurosci. 35, 9424–9431.

5. Arttamangkul, S., Platt, E.J., Carroll, J., Farrens, D., 2021. Functional independence of endogenous mu- and delta-opioid receptors co-expressed in cholinergic interneurons. Elife 10. 10.7554/eLife.69740.sa2

6. American Psychiatric Association, 2013. Diagnostic and statistical manual of mental disorders (DSM- 5®). American Psychiatric Pub.

7. Barr, C.S., Schwandt, M., Lindell, S.G., Chen, S.A., Goldman, D., Suomi, S.J., Higley, J.D., Heilig, M., 2007. Association of a functional polymorphism in the mu-opioid receptor gene with alcohol response and consumption in male rhesus macaques. Arch. Gen. Psychiatry 64, 369–376.

8. Barson, J.R., Carr, A.J., Soun, J.E., Sobhani, N.C., Leibowitz, S.F., Hoebel, B.G., 2009. Opioids in the nucleus accumbens stimulate ethanol intake. Physiol. Behav. 98, 453–459.

9. Bart, G., Kreek, M.J., Ott, J., LaForge, K.S., Proudnikov, D., Pollak, L., Heilig, M., 2005. Increased attributable risk related to a functional mu-opioid receptor gene polymorphism in association with alcohol dependence in central Sweden. Neuropsychopharmacology 30, 417–422.

10. Becker, A., Grecksch, G., Kraus, J., Loh, H.H., Schroeder, H., Höllt, V., 2002. Rewarding effects of ethanol and cocaine in µ opioid receptor-deficient mice. Naunyn-Schmiedeberg’s Arch Pharmacol 365, 296–302.

11. Berlanga, M.L., Olsen, C.M., Chen, V., Ikegami, A., Herring, B.E., Duvauchelle, C.L., Alcantara, A.A., 2003. Cholinergic interneurons of the nucleus accumbens and dorsal striatum are activated by the self-administration of cocaine. Neuroscience 120, 1149–1156.

12. Berrendero, F., Robledo, P., Trigo, J.M., Martín-García, E., Maldonado, R., 2010. Neurobiological mechanisms involved in nicotine dependence and reward: participation of the endogenous opioid system. Neurosci. Biobehav. Rev. 35, 220–231.

13. Blomqvist, O., Ericson, M., Johnson, D.H., Engel, J.A., Söderpalm, B., 1996. Voluntary ethanol intake in the rat: effects of nicotinic acetylcholine receptor blockade or subchronic nicotine treatment. Eur. J. Pharmacol. 314, 257–267.

14. Britt, J.P., McGehee, D.S., 2008. Presynaptic opioid and nicotinic receptor modulation of dopamine overflow in the nucleus accumbens. J. Neurosci. 28, 1672–1681.

15. Collins, A.C., Pogun, S., Nesil, T., Kanit, L., 2012. Oral Nicotine Self-Administration in Rodents. J. Addict. Res. Ther. S2. 10.4172/2155-6105.S2-004

16. Corrigall, W.A., Coen, K.M., Adamson, K.L., Chow, B.L., 1999. Manipulations of mu-opioid and nicotinic cholinergic receptors in the pontine tegmental region alter cocaine self-administration in rats. Psychopharmacology 145, 412–417.

17. Cui, Y., Ostlund, S.B., James, A.S., Park, C.S., Ge, W., Roberts, K.W., Mittal, N., Murphy, N.P., Cepeda, C., Kieffer, B.L., Levine, M.S., Jentsch, J.D., Walwyn, W.M., Sun, Y.E., Evans, C.J., Maidment, N.T., Yang, X.W., 2014. Targeted expression of μ-opioid receptors in a subset of striatal direct-pathway neurons restores opiate reward. Nat. Neurosci. 17, 254–261.

18. DeBaker, M.C., Robinson, J.M., Moen, J.K., Wickman, K., Lee, A.M., 2020. Differential patterns of alcohol and nicotine intake: Combined alcohol and nicotine binge consumption behaviors in mice. Alcohol 85, 57–64.

19. Dhatt, R.K., Gudehithlu, K.P., Wemlinger, T.A., Tejwani, G.A., Neff, N.H., Hadjiconstantinou, M., 1995. Preproenkephalin mRNA and methionine-enkephalin content are increased in mouse striatum after treatment with nicotine. J. Neurochem. 64, 1878–1883.

20. Erbs, E., Faget, L., Scherrer, G., Matifas, A., Filliol, D., Vonesch, J.-L., Koch, M., Kessler, P., Hentsch, D., Birling, M.-C., Koutsourakis, M., Vasseur, L., Veinante, P., Kieffer, B.L., Massotte, D., 2015. A mu-delta opioid receptor brain atlas reveals neuronal co-occurrence in subcortical networks. Brain Struct. Funct. 220, 677–702.

21. Ericson, M., Löf, E., Stomberg, R., Söderpalm, B., 2009. The smoking cessation medication varenicline attenuates alcohol and nicotine interactions in the rat mesolimbic dopamine system. J. Pharmacol. Exp. Ther. 329, 225–230.

22. Feduccia, A.A., Simms, J.A., Mill, D., Yi, H.Y., Bartlett, S.E., 2014. Varenicline decreases ethanol intake and increases dopamine release via neuronal nicotinic acetylcholine receptors in the nucleus accumbens. Br. J. Pharmacol. 171, 3420–3431.

23. Galaj, E., Kipp, B.T., Floresco, S.B., Savage, L.M., 2019. Persistent Alterations of Accumbal Cholinergic Interneurons and Cognitive Dysfunction after Adolescent Intermittent Ethanol Exposure. Neuroscience 404, 153–164.

24. Ghozland, S., Chu, K., Kieffer, B.L., Roberts, A.J., 2005. Lack of stimulant and anxiolytic-like effects of ethanol and accelerated development of ethanol dependence in mu-opioid receptor knockout mice. Neuropharmacology 49, 493–501.

25. Gianoulakis, C., 2009. Endogenous opioids and addiction to alcohol and other drugs of abuse. Curr. Top. Med. Chem. 9, 999–1015.

26. Gilpin, N.W., Richardson, H.N., Koob, G.F., 2008. Effects of CRF1-receptor and opioid-receptor antagonists on dependence-induced increases in alcohol drinking by alcohol-preferring (P) rats. Alcohol. Clin. Exp. Res. 32, 1535–1542.

27. Gonzales, K.K., Smith, Y., 2015. Cholinergic interneurons in the dorsal and ventral striatum: anatomical and functional considerations in normal and diseased conditions. Ann. N. Y. Acad. Sci. 1349, 1–45.

28. Hall, F.S., Sora, I., Uhl, G.R., 2001. Ethanol consumption and reward are decreased in mu-opiate receptor knockout mice. Psychopharmacology 154, 43–49.

29. Heinz, A., Reimold, M., Wrase, J., Hermann, D., Croissant, B., Mundle, G., Dohmen, B.M., Braus, D.F., Schumann, G., Machulla, H.-J., Bares, R., Mann, K., 2005. Correlation of stable elevations in striatal mu-opioid receptor availability in detoxified alcoholic patients with alcohol craving: a positron emission tomography study using carbon 11-labeled carfentanil. Arch. Gen. Psychiatry 62, 57–64.

30. Hendrickson, L.M., Guildford, M.J., Tapper, A.R., 2013. Neuronal nicotinic acetylcholine receptors: common molecular substrates of nicotine and alcohol dependence. Front. Psychiatry 4, 29.

31. Hendrickson, L.M., Zhao-Shea, R., Pang, X., Gardner, P.D., Tapper, A.R., 2010. Activation of alpha4* nAChRs is necessary and sufficient for varenicline-induced reduction of alcohol consumption. J. Neurosci. 30, 10169–10176.

32. Hermann, D., Hirth, N., Reimold, M., Batra, A., Smolka, M.N., Hoffmann, S., Kiefer, F., Noori, H.R., Sommer, W.H., Reischl, G., la Fougère, C., Mann, K., Spanagel, R., Hansson, A.C., 2017. Low μ- Opioid Receptor Status in Alcohol Dependence Identified by Combined Positron Emission Tomography and Post-Mortem Brain Analysis. Neuropsychopharmacology 42, 606–614.

33. Herring, B.E., Mayfield, R.D., Camp, M.C., Alcantara, A.A., 2004. Ethanol-induced Fos immunoreactivity in the extended amygdala and hypothalamus of the rat brain: focus on cholinergic interneurons of the nucleus accumbens. Alcohol. Clin. Exp. Res. 28, 588–597.

34. Heyser, C.J., Roberts, A.J., Schulteis, G., Koob, G.F., 1999. Central administration of an opiate antagonist decreases oral ethanol self-administration in rats. Alcohol. Clin. Exp. Res. 23, 1468–1476.

35. Hopf, F.W., Lesscher, H.M.B., 2014. Rodent models for compulsive alcohol intake. Alcohol 48, 253– 264.

36. Houdi, A.A., Pierzchala, K., Marson, L., Palkovits, M., Van Loon, G.R., 1991. Nicotine-induced alteration in Tyr-Gly-Gly and Met-enkephalin in discrete brain nuclei reflects altered enkephalin neuron activity. Peptides 12, 161–166.

37. Hyytiä, P., Kiianmaa, K., 2001. Suppression of ethanol responding by centrally administered CTOP and naltrindole in AA and Wistar rats. Alcohol. Clin. Exp. Res. 25, 25–33.

38. Jabourian, M., Venance, L., Bourgoin, S., Ozon, S., Pérez, S., Godeheu, G., Glowinski, J., Kemel, M.-L., 2005. Functional mu opioid receptors are expressed in cholinergic interneurons of the rat dorsal striatum: territorial specificity and diurnal variation. Eur. J. Neurosci. 21, 3301–3309.

39. Kiguchi, Y., Aono, Y., Watanabe, Y., Yamamoto-Nemoto, S., Shimizu, K., Shimizu, T., Kosuge, Y., Waddington, J.L., Ishige, K., Ito, Y., Saigusa, T., 2016. In vivo neurochemical evidence that delta1-, delta2- and mu2-opioid receptors, but not mu1-opioid receptors, inhibit acetylcholine efflux in the nucleus accumbens of freely moving rats. Eur. J. Pharmacol. 789, 402–410.

40. Kolpakova, J., van der Vinne, V., Gimenez-Gomez, P., Le, T., Martin, G.E., 2022. Binge alcohol drinking alters the differential control of cholinergic interneurons over nucleus accumbens D1 and D2 medium spiny neurons. bioRxiv. 10.1101/2022.06.22.497229

41. Lam, M.P., Nurmi, H., Rouvinen, N., Kiianmaa, K., Gianoulakis, C., 2010. Effects of acute ethanol on beta-endorphin release in the nucleus accumbens of selectively bred lines of alcohol-preferring AA and alcohol-avoiding ANA rats. Psychopharmacology 208, 121–130.

42. Learn, J.E., Chernet, E., McBride, W.J., Lumeng, L., Li, T.-K., 2001. Quantitative Autoradiography of Mu-Opioid Receptors in the CNS of High--Alcohol-Drinking (HAD) and Low--Alcohol-Drinking (LAD) Rats. Alcohol. Clin. Exp. Res. 25, 524–530.

43. Le Moine, C., Kieffer, B., Gaveriaux-Ruff, C., Befort, K., Bloch, B., 1994. Delta-opioid receptor gene expression in the mouse forebrain: localization in cholinergic neurons of the striatum. Neuroscience 62, 635–640.

44. Lesscher, H.M.B., van Kerkhof, L.W.M., Vanderschuren, L.J.M.J., 2010. Inflexible and indifferent alcohol drinking in male mice. Alcohol. Clin. Exp. Res. 34, 1219–1225.

45. Leyrer-Jackson, J.M., Holter, M., Overby, P.F., Newbern, J.M., Scofield, M.D., Olive, M.F., Gipson, C.D., 2021. Accumbens Cholinergic Interneurons Mediate Cue-Induced Nicotine Seeking and Associated Glutamatergic Plasticity. eNeuro 8. 10.1523/ENEURO.0276-20.2020

46. Liu, C., Cai, X., Ritzau-Jost, A., Kramer, P.F., Li, Y., Khaliq, Z.M., Hallermann, S., Kaeser, P.S., 2022. An action potential initiation mechanism in distal axons for the control of dopamine release. Science 375, 1378–1385.

47. Loftén, A., Adermark, L., Ericson, M., Söderpalm, B., 2023. Regulation of ethanol_mediated dopamine elevation by glycine receptors located on cholinergic interneurons in the nucleus accumbens. Addict. Biol. 28. 10.1111/adb.13349

48. Loftén, A., Adermark, L., Ericson, M., Söderpalm, B., 2021. An acetylcholine-dopamine interaction in the nucleus accumbens and its involvement in ethanol’s dopamine-releasing effect. Addict. Biol. 26, e12959.

49. Mansvelder, H.D., De Rover, M., McGehee, D.S., Brussaard, A.B., 2003. Cholinergic modulation of dopaminergic reward areas: upstream and downstream targets of nicotine addiction. Eur. J. Pharmacol. 480, 117–123.

50. Marty, M.A., Erwin, V.G., Cornell, K., Zgombick, J.M., 1985. Effects of nicotine on β-endorphin, αMSH, and ACTH secretion by isolated perfused mouse brains and pituitary glands, in Vitro. Pharmacol. Biochem. Behav. 22, 317–325.

51. Ma, T., Huang, Z., Xie, X., Cheng, Y., Zhuang, X., Childs, M.J., Gangal, H., Wang, X., Smith, L.N., Smith, R.J., Zhou, Y., Wang, J., 2022. Chronic alcohol drinking persistently suppresses thalamostriatal excitation of cholinergic neurons to impair cognitive flexibility. J. Clin. Invest. 132. 10.1172/JCI154969

52. Méndez, M., Barbosa-Luna, I.G., Pérez-Luna, J.M., Cupo, A., Oikawa, J., 2010. Effects of acute ethanol administration on methionine-enkephalin expression and release in regions of the rat brain. Neuropeptides 44, 413–420.

53. Méndez, M., Morales-Mulia, M., 2008. Role of mu and delta opioid receptors in alcohol drinking behaviour. Curr. Drug Abuse Rev. 1, 239–252.

54. Mitchell, J.M., O’Neil, J.P., Janabi, M., Marks, S.M., Jagust, W.J., Fields, H.L., 2012. Alcohol consumption induces endogenous opioid release in the human orbitofrontal cortex and nucleus accumbens. Sci. Transl. Med. 4, 116ra6.

55. Mohebi, A., Collins, V.L., Berke, J.D., 2023. Accumbens cholinergic interneurons dynamically promote dopamine release and enable motivation. Elife 12. 10.7554/eLife.85011

56. Nadal, R., Chappell, A.M., Samson, H.H., 1998. Effects of nicotine and mecamylamine microinjections into the nucleus accumbens on ethanol and sucrose self_administration. Alcohol. Clin. Exp. Res. 22, 1190–1198.

57. Nealey, K.A., Smith, A.W., Davis, S.M., Smith, D.G., Walker, B.M., 2011. κ-opioid receptors are implicated in the increased potency of intra-accumbens nalmefene in ethanol-dependent rats. Neuropharmacology 61, 35–42.

58. Okada, K., Nishizawa, K., Setogawa, S., Hashimoto, K., Kobayashi, K., 2018. Task-dependent function of striatal cholinergic interneurons in behavioural flexibility. Eur. J. Neurosci. 47, 1174–1183.

59. Olive, M.F., Koenig, H.N., Nannini, M.A., Hodge, C.W., 2001. Stimulation of endorphin neurotransmission in the nucleus accumbens by ethanol, cocaine, and amphetamine. J. Neurosci. 21, RC184.

60. Oslin, D.W., Berrettini, W., Kranzler, H.R., Pettinati, H., Gelernter, J., Volpicelli, J.R., O’Brien, C.P., 2003. A functional polymorphism of the mu-opioid receptor gene is associated with naltrexone response in alcohol-dependent patients. Neuropsychopharmacology 28, 1546–1552.

61. Oude Ophuis, R.J.A., Boender, A.J., van Rozen, A.J., Adan, R.A.H., 2014. Cannabinoid, melanocortin and opioid receptor expression on DRD1 and DRD2 subpopulations in rat striatum. Front. Neuroanat. 8, 14.

62. Perry, C.J., McNally, G.P., 2013. μ-Opioid receptors in the nucleus accumbens shell mediate context-induced reinstatement (renewal) but not primed reinstatement of extinguished alcohol seeking. Behav. Neurosci. 127, 535–543.

63. Perry, T.W., Sneddon, E.A., Reichert, A.N., Schuh, K.M., Shand, N.A., Quinn, J.J., Radke, A.K., 2023. Sex, but not early life stress, effects on two-bottle choice alcohol drinking behaviors in mice. bioRxiv. 10.1101/2023.01.21.524642

64. Pogun, S., Yararbas, G., Nesil, T., Kanit, L., 2017. Sex differences in nicotine preference. J. Neurosci. Res. 95, 148–162.

65. Ponterio, G., Tassone, A., Sciamanna, G., Riahi, E., Vanni, V., Bonsi, P., Pisani, A., 2013. Powerful inhibitory action of mu opioid receptors (MOR) on cholinergic interneuron excitability in the dorsal striatum. Neuropharmacology 75, 78–85.

66. Rada, P., Jensen, K., Hoebel, B.G., 2001. Effects of nicotine and mecamylamine-induced withdrawal on extracellular dopamine and acetylcholine in the rat nucleus accumbens. Psychopharmacology 157, 105–110.

67. Radke, A.K., Sneddon, E.A., Frasier, R.M., Hopf, F.W., 2021. Recent Perspectives on Sex Differences in Compulsion-Like and Binge Alcohol Drinking. Int. J. Mol. Sci. 22, 3788.

68. Radke, A.K., Held, I.T., Sneddon, E.A., Riddle, C.A., Quinn, J.J., 2020. Additive influences of acute early life stress and sex on vulnerability for aversion-resistant alcohol drinking. Addict. Biol. 25, e12829.

69. Richard, J.M., Fields, H.L., 2016. Mu-opioid receptor activation in the medial shell of nucleus accumbens promotes alcohol consumption, self-administration and cue-induced reinstatement. Neuropharmacology 108, 14–23.

70. Roberts, A.J., McDonald, J.S., Heyser, C.J., Kieffer, B.L., Matthes, H.W.D., Koob, G.F., Gold, L.H., 2000. μ-Opioid Receptor Knockout Mice Do Not Self-Administer Alcohol ,. J. Pharmacol. Exp. Ther. 293, 1002–1008.

71. Schilaty, N.D., Hedges, D.M., Jang, E.Y., Folsom, R.J., Yorgason, J.T., McIntosh, J.M., Steffensen, S.C., 2014. Acute ethanol inhibits dopamine release in the nucleus accumbens via α6 nicotinic acetylcholine receptors. J. Pharmacol. Exp. Ther. 349, 559–567.

72. Schuh, K.M., Sneddon, E.A., Nader, A.M., Muench, M.A., Radke, A.K., 2022. Orbitofrontal cortex subregion inhibition during binge-like and aversion-resistant alcohol drinking. Alcohol 99, 1–8.

73. Seif, T., Chang, S.-J., Simms, J.A., Gibb, S.L., Dadgar, J., Chen, B.T., Harvey, B.K., Ron, D., Messing, R.O., Bonci, A., Hopf, F.W., 2013. Cortical activation of accumbens hyperpolarization-active NMDARs mediates aversion-resistant alcohol intake. Nat. Neurosci. 16, 1094–1100.

74. Serafin, M., Khateb, A., Mühlethaler, M., 1990. Opiates inhibit pedunculopontine neurones in guinea pig brainstem slices. Neurosci. Lett. 119, 125–128.

75. Severino, A.L., Mittal, N., Hakimian, J.K., Velarde, N., Minasyan, A., Albert, R., Torres, C., Romaneschi, N., Johnston, C., Tiwari, S., Lee, A.S., Taylor, A.M., Gavériaux-Ruff, C., Kieffer, B.L., Evans, C.J., Cahill, C.M., Walwyn, W.M., 2020. μ-Opioid Receptors on Distinct Neuronal Populations Mediate Different Aspects of Opioid Reward-Related Behaviors. eNeuro 7. 10.1523/ENEURO.0146-20.2020

76. Sharma, R., Chischolm, A., Parikh, M., Thakkar, M., 2024. Cholinergic interneurons in the shell region of the nucleus accumbens regulate binge alcohol consumption: A chemogenetic and genetic lesion study. Alcohol. Clin. Exp. Res. 10.1111/acer.15295.

77. Sneddon, E.A., Schuh, K.M., Frankel, J.W., Radke, A.K., 2021. The contribution of medium spiny neuron subtypes in the nucleus accumbens core to compulsive-like ethanol drinking. Neuropharmacology 187, 108497.

78. Sneddon, E.A., White, R.D., Radke, A.K., 2019. Sex Differences in Binge-Like and Aversion-Resistant Alcohol Drinking in C57BL/6J Mice. Alcohol. Clin. Exp. Res. 43, 243–249.

79. Sneddon, E.A., Schuh, K.M., Frankel, J.W., Radke, A.K., 2021. The contribution of medium spiny neuron subtypes in the nucleus accumbens core to compulsive-like ethanol drinking. Neuropharmacology 187, 108497.

80. Sprow, G.M., Thiele, T.E., 2012. The neurobiology of binge-like ethanol drinking: evidence from rodent models. Physiol. Behav. 106, 325–331.

81. Steensland, P., Simms, J.A., Holgate, J., Richards, J.K., Bartlett, S.E., 2007. Varenicline, an α4β2 nicotinic acetylcholine receptor partial agonist, selectively decreases ethanol consumption and seeking. Proceedings of the National Academy of Sciences 104, 12518–12523.

82. Steidl, S., Wasserman, D.I., Blaha, C.D., Yeomans, J.S., 2017. Opioid-induced rewards, locomotion, and dopamine activation: A proposed model for control by mesopontine and rostromedial tegmental neurons. Neurosci. Biobehav. Rev. 83, 72–82.

83. Stowe, T.A., Pitts, E.G., Leach, A.C., Iacino, M.C., Niere, F., Graul, B., Raab-Graham, K.F., Yorgason, J.T., Ferris, M.J., 2022. Diurnal rhythms in cholinergic modulation of rapid dopamine signals and associative learning in the striatum. Cell Rep. 39, 110633.

84. Trigo, J.M., Zimmer, A., Maldonado, R., 2009. Nicotine anxiogenic and rewarding effects are decreased in mice lacking beta-endorphin. Neuropharmacology 56, 1147–1153.

85. Wadsworth, H.A., Anderson, E.Q., Williams, B.M., Ronström, J.W., Moen, J.K., Lee, A.M., McIntosh, J.M., Wu, J., Yorgason, J.T., Steffensen, S.C., 2023. Role of α6-Nicotinic Receptors in Alcohol-Induced GABAergic Synaptic Transmission and Plasticity to Cholinergic Interneurons in the Nucleus Accumbens. Mol. Neurobiol. 60, 3113–3129.

86. Walters, C.L., Cleck, J.N., Kuo, Y.-C., Blendy, J.A., 2005. Mu-opioid receptor and CREB activation are required for nicotine reward. Neuron 46, 933–943.

87. Weinberger, A.H., Pacek, L.R., Giovenco, D., Galea, S., Zvolensky, M.J., Gbedemah, M., Goodwin, R.D., 2019. Cigarette Use Among Individuals with Alcohol Use Disorders in the United States, 2002 to 2016: Trends Overall and by Race/Ethnicity. Alcohol. Clin. Exp. Res. 43, 79–90.

88. Williams, E.A., Bradley, C.A., Palmatier, M.I., 2017. Silencing giant cholinergic interneurons in the nucleus accumbens with DREADDS reduces nicotine self-administration in rats. Drug & Alcohol Dependence 171, e217–e218.

89. Wouda, J.A., Riga, D., De Vries, W., Stegeman, M., van Mourik, Y., Schetters, D., Schoffelmeer, A.N.M., Pattij, T., De Vries, T.J., 2011. Varenicline attenuates cue-induced relapse to alcohol, but not nicotine seeking, while reducing inhibitory response control. Psychopharmacology 216, 267–277.

90. Yoo, J.-H., Lee, S.-Y., Loh, H.H., Ho, I.K., Jang, C.-G., 2004. Loss of nicotine-induced behavioral sensitization in micro-opioid receptor knockout mice. Synapse 51, 219–223.

91. Yorgason, J.T., Zeppenfeld, D.M., Williams, J.T., 2017. Cholinergic Interneurons Underlie Spontaneous Dopamine Release in Nucleus Accumbens. J. Neurosci. 37, 2086–2096.

92. Zhang, M., Kelley, A.E., 2002. Intake of saccharin, salt, and ethanol solutions is increased by infusion of a mu opioid agonist into the nucleus accumbens. Psychopharmacology 159, 415–423.

